# The Derived-Band Envelope Following Response and its Sensitivity to Sensorineural Hearing Deficits

**DOI:** 10.1101/820704

**Authors:** Sarineh Keshishzadeh, Markus Garrett, Viacheslav Vasilkov, Sarah Verhulst

## Abstract

The envelope following response (EFR) has been proposed as a non-invasive marker of synaptopathy in animal models. However, its amplitude is affected by the spread of basilar-membrane excitation and other coexisting sensorineural hearing deficits. This study aims to (i) improve frequency specificity of the EFR by introducing a derived-band EFR (DBEFR) technique and (ii) investigate the effect of lifetime noise exposure, age and outer-hair-cell (OHC) damage on DBEFR magnitudes. Additionally, we adopt a modelling approach to validate the frequency-specificity of the DBEFR and test how different aspects of sensorineural hearing loss affect peripheral generators. The combined analysis of simulations and experimental data proposes that the DBEFRs extracted from the [2-6]-kHz frequency band is a sensitive and frequency-specific measure of synaptopathy in humans. Individual variability in DBEFR magnitudes among listeners with normal audiograms was explained by their self-reported amount of experienced lifetime noise-exposure and corresponded to amplitude variability predicted by synaptopathy. Older listeners consistently had reduced DBEFR magnitudes in comparison to young normal-hearing listeners, in correspondence to how age-induced synaptopathy affects EFRs and compromises temporal envelope encoding. Lastly, OHC damage was also seen to affect the DBEFR magnitude, hence this marker should be combined with a sensitive marker of OHC-damage to offer a differential diagnosis of synaptopathy in listeners with impaired audiograms.

## 1. Introduction

Struggling to understand speech in noisy environments is a prevalent complaint of the ageing population, even when they have normal audiometric thresholds. Although hearing thresholds are informative about the sensory function of the cochlea, they are insensitive to auditory-nerve (AN) fiber loss, which is the first sign of permanent hearing damage (Kujawa and Liberman, 2009; Liberman and Kujawa, 2017) and related to supra-threshold hearing (Bharadwaj et al., 2014). Recent animal studies have shown that ageing, ototoxicity and overexposure to noise can lead to an irreversible loss of AN synapses, i.e. cochlear synaptopathy (CS), and delayed degeneration of cochlear neurons, while leaving the cochlear sensory hair cells intact (Kujawa and Liberman, 2009; Lin et al., 2011; Liu et al., 2012; Furman et al., 2013; Lobarinas et al., 2017; Valero et al., 2017). Even when the noise exposure dose only causes a temporary threshold shift (Kujawa and Liberman, 2009), noise-induced AN fibers degeneration can progress through the lifespan and yield an increased sensitivity of the ear to age-induced hearing dysfunction (Fernandez et al., 2015). Additionally, reduced numbers of spiral ganglion cells in post-mortem histology of human temporal bones with preserved sensory cells, confirmed the existence of age-related CS in humans (Makary et al., 2011; Viana et al., 2015; Wu et al., 2019). Thus, noise exposure and ageing are important causes of CS, a deficit which compromises the temporal coding fidelity of supra-threshold sound as a result of a reduced number of afferent AN synapses innervating the inner hair cell (Bharadwaj et al., 2014, 2015).

Since the discovery of CS, several attempts have been made to associate changes in indirect and non-invasive measures of auditory function such as scalp-recorded auditory evoked potentials (AEPs) to the histologically quantified degree of AN fibers loss in animals. For example, auditory brainstem responses (ABRs), evoked by transient stimuli and reflecting the synchronized onset responses of AN fibers (Don and Eggermont, 1978) showed a decreased supra-threshold wave-I amplitude after synaptopathy due to noise-exposure (Kujawa and Liberman, 2009; Lobarinas et al., 2017; Lin et al., 2011), despite recovered normal distortion product otoacoustic emission (DPOAE) and ABR thresholds. The number of AN fibers can also be quantified using envelope following responses (EFRs), which capture how well AN fibers can phase-lock to the stimulus envelope (Joris and Yin, 1992). The EFR can be extracted from scalp-electrodes in response to a sinusoidally amplitude modulated (SAM) pure-tone stimulus (Bharadwaj et al., 2014), and has been proposed as an AEP-based measure of CS (Shaheen et al., 2015; Parthasarathy and Kujawa, 2018).

Despite the strong relation between AEP markers and CS in animal studies, the indirect nature of AEP recordings hinders a clear and direct interpretation of response strength in terms of CS. First of all, a mixture of sources contribute to scalp potentials, some of which are electrical activity induced by subject-specific factors and unrelated to the sound-driven response (e.g. head size, age, sex, geometry of the generators and physiological noise level; Trune et al., 1988; Mitchell et al., 1989; Bharadwaj et al., 2014; Plack et al., 2016). Other sources relate to the sound-driven response but depend on outer-hair-cell (OHC) health (Gorga et al., 1985) or cochlear tonotopy (Don and Eggermont, 1978). Lastly, the scalp-recorded AEP is strongly influenced by stimulus characteristics and the corresponding spread of basilar-membrane (BM) excitation, which can confound a frequency-specific diagnosis of CS (Bharadwaj et al., 2014, 2015; Verhulst et al., 2018a; Encina-Llamas et al., 2019). To address these issues, several studies have proposed differential/relative AEP-based metrics: the EFR amplitude slope as a function of modulation depth (Bharadwaj et al., 2014, 2015; Guest et al., 2018), ABR wave-V latency changes in different levels of background noise (Mehraei et al., 2016), or the combined use of noise-floor corrected EFRs with ABRs to segregate mixed hearing pathologies and normalize inter-individual variabilities (Vasilkov and Verhulst, 2019, preprint). Secondly, a number of techniques have been proposed to confine ABR generation to specific frequency bands: the use of simultaneous off-frequency masking paradigms, i.e. the derived-band ABR (Eggermont, 1976; Don and Eggermont, 1978), tone-burst ABRs (Rasetshwane et al., 2013) and notched noise paradigms (Abdala and Folsom, 1995). Lastly, asynchrony of low-spontaneous rate (LSR) AN fibers to the transient stimulus (Bourien et al., 2014) may limit the use of the ABR wave-I amplitude to capture all aspects of CS, as noise-induced CS might preferentially affect LSR AN fibers (Furman et al., 2013).

This study proposes the use of a relative derived-band EFR method (DBEFR), to confine the EFR to a specific frequency band. To construct DBEFRs, we changed the bandwidth of the stimulus on the low-frequency side rather than using off-frequency masking methods. Thus, a consecutive subtraction of responses to stimuli with various bandwidths will yield a relative measure of supra-threshold sound coding. We further hypothesize that the relative metric design of the DBEFR reduces the impact of subject-specific factors and increases its sensitivity to individual sensorineural hearing deficits. DBEFR magnitudes were extracted from individuals in four groups to study their applicability to diagnose sensorineural hearing deficits: (1) a young normal-hearing control group, (2) a group with self-reported hearing difficulties in noisy environments, (3) a group of older listeners with normal audiograms and (4) an age-matched group with sloping high-frequency audiograms. We assumed that the second group might be affected by CS due to noise overexposure or ageing and that the third group might be affected by age-induced CS, without co-occuring OHC damage. Aside from collecting DBEFRs, we assessed individual OHC function using audiometric and DPOAE thresholds. In line with animal studies of age-related and noise-induced synaptopathy, we expect that the DBEFR will be reduced in all but the control group.

Because, a direct assessment of the individual degree of OHC and AN damage is presently experimentally impossible, we complemented our experimental work with a modelling approach to better understand the relationship between sensorineural pathologies and their effect on the peripheral generators of the DBEFR. Models can study how AN fiber and sensory hair cell damage impacts the EFR generators to understand their respective roles for DBEFR generation (Verhulst et al., 2016, 2018a,b). We adopt a biophysically inspired model of the human auditory periphery calibrated for ABR and EFR simulation (Verhulst et al., 2018a) and considered the simulations together with the data to interpret the implications of our findings for DBEFR-based hearing diagnostics.

## 2. Materials and Methods

Two experiments were conducted at two recording locations. In the first experiment (Ghent University), normal-hearing (NH) and listeners with self-reported hearing difficulties (NHSR) participated. In the second experiment (Oldenburg University), a total of 43 participants were recruited in three groups: a young NH control group (yNH), an older NH group (oNH) and an older group with sloping high-frequency audiogram (oHI). Ethical approvals were obtained from Ghent and Oldenburg Universities and all participants were informed about the experimental procedures and signed an informed consent before the experiment.

### 2.1. Participants

16 NH listeners with ages between 18 and 30 (NH: 24.21±4.10 years, five females) and 9 NH subjects with self-reported hearing difficulties (NHSR) with ages between 23 to 49 (NHSR: 33.78±8.57 years, three females) participated in the first experiment. The NHSR participants were recruited using a flyer asking whether they had speech understanding difficulties in the presence of background noise, while not presently being treated for hearing disorders. Measurements were conducted in two sessions per subject, with a maximum sound exposure time of 90 minutes per session. The participants filled out a questionnaire, in which they were asked how often (yearly, monthly, weekly or daily) they had been playing a musical instrument in a band, attended festivals, concerts or discotheques and used noisy tools during their lifetime. Moreover, the total number of noise-exposed sessions, their duration and estimated noise loudness (a score between 1 to 5) were also assessed (Degeest et al., 2014). Audiograms were measured with an Interacoustics Clinical Computer Audiometer (AC5) at ten standard frequencies between 0.25 and 8 kHz.

The second experiment was conducted with three participant groups composed of: 15 young normal-hearing (yNH: 24.53±2.26 years, eight female), 16 old normal-hearing (oNH: 64.25±1.88 years, eight female) and 12 old hearing-impaired (oHI: 65.33±1.87 years, seven female) participants. All yNH participants had puretone thresholds below 20 dB-HL at all measured frequencies between 0.125 and 10 kHz (Auritec AT900, Hamburg, Germany audiometer). In both experiments, the audiometrically better ear was chosen for the experiment and stimuli were presented monaurally while participants were seated in a comfortable chair in an acoustically and electrically shielded sound booth, watching silent movies with subtitles to stay awake. Figure 1 shows audiograms of the subjects in all groups. From here on, Δ stands for the NH group in the first experiment, □ for NHSR group, ◊ for yNH in the second experiment, *○* for oNH and ◃ for oHI group.

**Figure 1.**
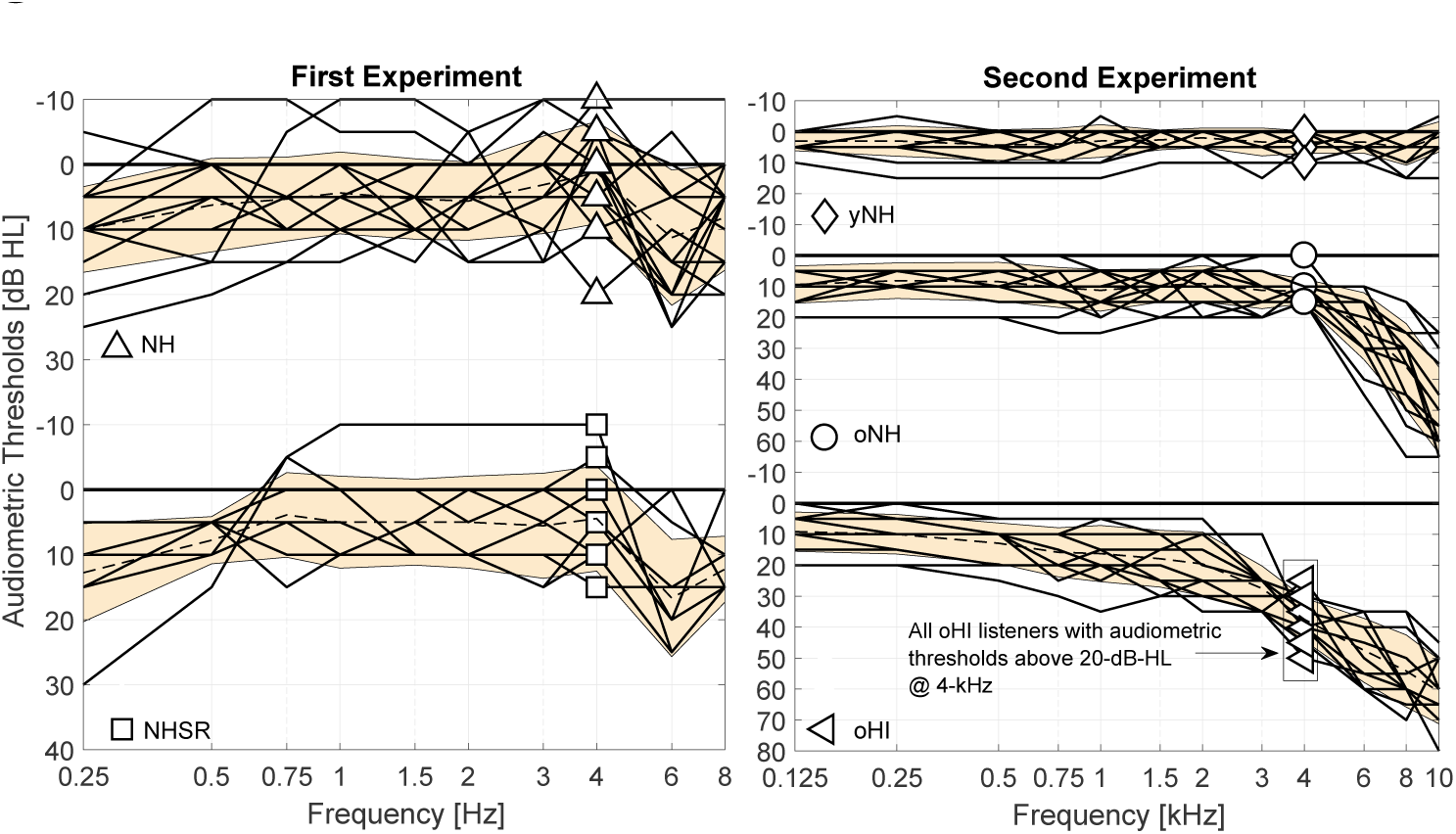
Measured audiograms in the first (left) and second (right) experiment. Markers indicate the audiometric threshold at 4 kHz. The dashed line is the averaged audiometric threshold at each group and the yellow shading the standard deviation.

### 2.2. Distortion Product Otoacoustic Emissions (DPOAEs)

In the first experiment, DPOAEs were recorded to ten primary-level pairs, (L_1_, L_2_), at nine primary-frequency pairs: f_2_ =[546, 780, 1002, 1476, 1998, 3012, 3996, 6006, 8003] and f_1_ = f_2_*/*1.2. L_2_ ranged from 20 to 65 dB-SPL in 5 dB steps and L_1_ = 0.4L_2_ + 39 dB, according to the scissors paradigm (Kummer et al., 1998). The nine primary frequency pairs were chosen to have complete stimulus periods of the primaries in each pair. For each frequency and level pair, 45 repetitions were generated in MATLAB 2016b and an ER-10X extended-bandwidth Etymotic Research probe system was used to deliver the two pure tones via a loudspeaker/microphone probe inserted in the ear-canal using a silicone eartip. The response was recorded and digitized using a Fireface UCX external sound card (RME). The pure tones were calibrated separately using a B&K artificial ear and B&K sound level meter at each primary frequency, separately. The time-domain ear-canal recordings were converted to pressure using the microphone sensitivity 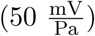 and pre-amplifier gain (40 dB). Then, I/O functions were calculated for the measured primary-frequency pairs by defining the L_DP_ as the averaged spectrum magnitude at the 2f_1_-f_2_ cubic distortion frequency, multiplied by 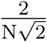, where N is the number of samples at each f_2_ response. Finally, a linear function, i.e. L_DP_ = aL_2_ + b, was fit to the bootstrapped data-points and the crossing point with L_DP_=0 Pa was defined as the DPOAE threshold at the measured f_2_ frequency. DPOAEs in the second experiment were acquired using a custom-made software (Mauermann, 2013) which implements a primary frequency sweep method at a fixed f_2_*/*f_1_ of 1.2 (Long et al., 2008). The primary frequencies were swept across an 1/3 octave range around the f_2_ = 4 kHz geometric mean with a duration of 2s/octave. Primary levels were chosen according to the scissors paradigm (Kummer et al., 1998). DPOAE threshold at each frequency was calculated by fitting a linear function to the bootstrapped data-points and was extrapolated to cross L_DP_=0 Pa. Additional details on the experimental procedure can be found in Verhulst et al. (2016).

### 2.3. Envelope Following Responses (EFRs)

The EFR stimuli in the first experiment were five filtered white noise carriers, which were 100% modulated with a 120-Hz sinusoid. To generate them, the white noise was filtered between the following frequency regions: [0.25-22], [0.5-22], [1-22], [2-22] and [4-22] kHz, using a 1024th order FIR band-pass filter designed by the Blackman-window method. In each frequency band, a stimulus with a duration of 1.25 s was generated in MATLAB 2016b, windowed with a 1.25% cosine-tapered window and delivered monaurally over ER-2 earphones, connected to a Fireface UCX external sound card (RME) and a TDT-HB7 headphone driver. A uniformly-distributed random silence jitter was applied between consecutive epochs (200 ms±20 ms) of the 370 stimulus presentations. Stimuli with various bandwidths were calibrated to have the same spectral magnitude, i.e. the widest bandwidth stimulus was presented at 70-dB-SPL, while narrower bandwidth stimuli had lower sound pressure levels to preserve an equal spectral level in all conditions. The calibration was performed using a B&K sound-level-meter type 2606. Figure 2a illustrates the designed stimuli in the frequency domain. Scalp-recorded potentials were obtained with a 64-Channel Biosemi EEG recording system and a custom-built trigger box using a sampling frequency of 16384 Hz. The electrodes were placed according to the 10-20 standard, using highly conductive gel (Signa gel). The Common Mode Sense (CMD) and Driven Right Leg (DRL) electrodes were placed on top of the head. Six external channels were used as well, i.e. two earlobe electrodes as reference and the remaining electrodes were placed on the forehead and cheeks to record electrical activity induced by horizontal and vertical eye movements. All channels were re-referenced to the average of the two earlobe electrodes.

**Figure 2.**
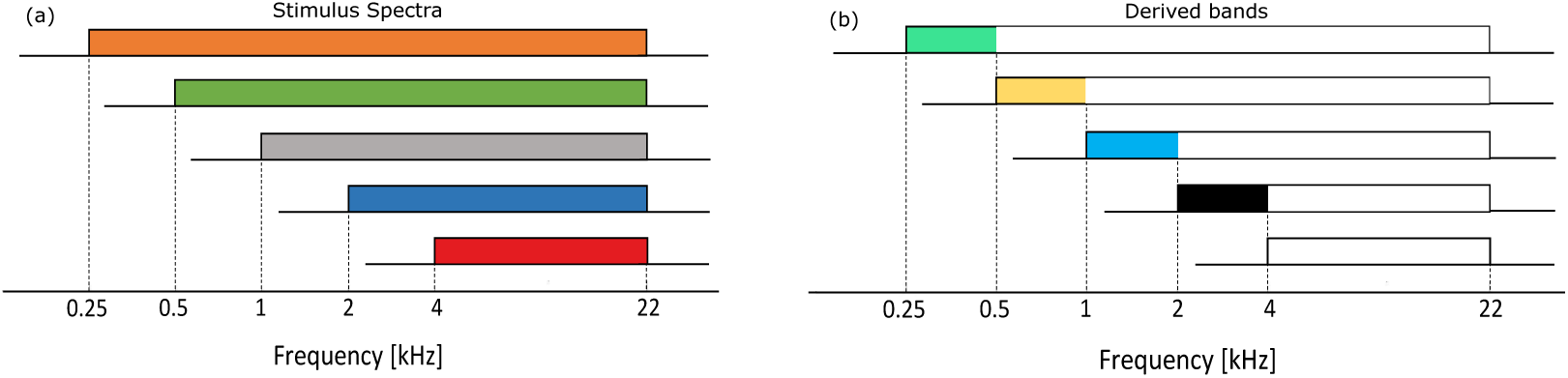
Spectra of the 120-Hz modulated stimuli and derived bands. (a) Designed stimulus spectra in different frequency bands and specified cut-off frequencies of the bandpass filter. (b) Derived bands from the EFRs recorded to the stimuli shown in (a) obtained by spectral subtraction.

In the second experiment, four EFR stimuli with white noise carriers were band-pass filtered using the same filter as in the first experiment in [0.3-16], [0.7-16], [2.8-16] and [5.6-16] kHz frequency regions. The precise lower cut-off frequencies employed in the band-pass filtering were 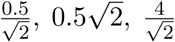 and 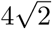 kHz, respectively. Stimuli were 95% modulated with a 120-Hz pure tone and presented at 70 dB SPL using the same configuration as the first experiment. The stimuli had a duration of 400 ms, were 2.5% ramped with a tapered-cosine window and presented 1000 times using a uniformly distributed random inter-stimulus silence jitter of 100 ms±10 ms. The calibration was performed in the same way as for the first experiment, but using B&K sound level meter type 2610. A 64-channel Biosemi EEG system was adopted to record the responses using EEG caps with equidistant electrode spacing. The CMS and DRL electrodes were located on the fronto-central midline and on the tip of the nose of the participants, respectively.

## 3. EFR Analysis

Acquired EFRs were first filtered using an 800th order Blackman window-based FIR filter between 60 and 600 Hz, using the *filtfilt* function of MATLAB to avoid time delays and phase shifts. Signals were broken into 1-s long epochs relative to the trigger onset, from 0.25 to 1.25 s in the first and into 0.3-s long epochs, from 0.1 to 0.4 s in the second experiment. Baseline correction was applied before the epochs were averaged across trials. 30 and 100 epochs were rejected on the basis of the highest peak-to-trough values in the first and second experiment, respectively. Since the firing patterns of neurons are influenced by factors such as instantaneous external inputs, previous firing patterns and the general state of the system, the interpretation of the raw EFR spectrum resulting from the Fast Fourier Transform (FFT) of the averaged epochs is challenging. Synaptic delays and axon conduction limitations cause a 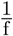 behaviour in EEG (Buzsaki, 2006, Chapter 10) and it is crucial to suppress this noise-floor to analyse the stimulus-driven spectrum. The bootstrapping approach proposed in Zhu et al. (2013) was employed to estimate the 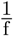 noise-floor component. First, 340 epochs were drawn randomly with replacement, among the 340 epochs (900 epochs in the second experiment). Then, the FFT of these epochs were averaged. This procedure was repeated N_1_=200 times (N_2_=400 for the second experiment), resulting in a nearly Gaussian distribution of raw, averaged spectra. The average value of this distribution yielded the frequency domain representation of the EFRs. Afterwards, the same procedure with M_1_=1000 repetitions (M_2_=1200 for the second experiment) and phase-flipped (180°) odd epochs was followed to estimate the spectral noise-floor as a function of frequency. The idea behind this approach is that the time-locked response is suppressed if the averaging is repeated sufficiently across phase-inverted epochs. Finally, the averaged absolute values of the estimated noise floors were subtracted from the averaged absolute values of the EFR spectra amplitudes to obtain the stimulus-driven EFR spectrum: 

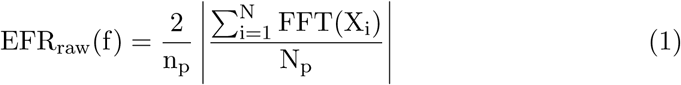

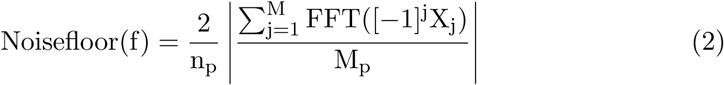

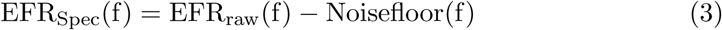

X represents the epochs vector, N the number of bootstrap repetitions, M the number of repetitions to estimate the noise-floor, p the experiment number (i.e. one or two) and n equals the number of FFT points (n_1_=16384 and n_2_=8192). Figure 3 represents EFR_raw_, Noisefloor and EFR_Spec_ spectra of subject No. 8 from NH group in the first experiment. All EFR_Spec_ peak values which were four standard deviations above the noise-floor (EFR_SpecSD_) for frequencies corresponding to the modulation frequency (120 Hz) and its following two harmonics (240 and 360 Hz) were added to yield EFR magnitude of the corresponding condition.

**Figure 3.**
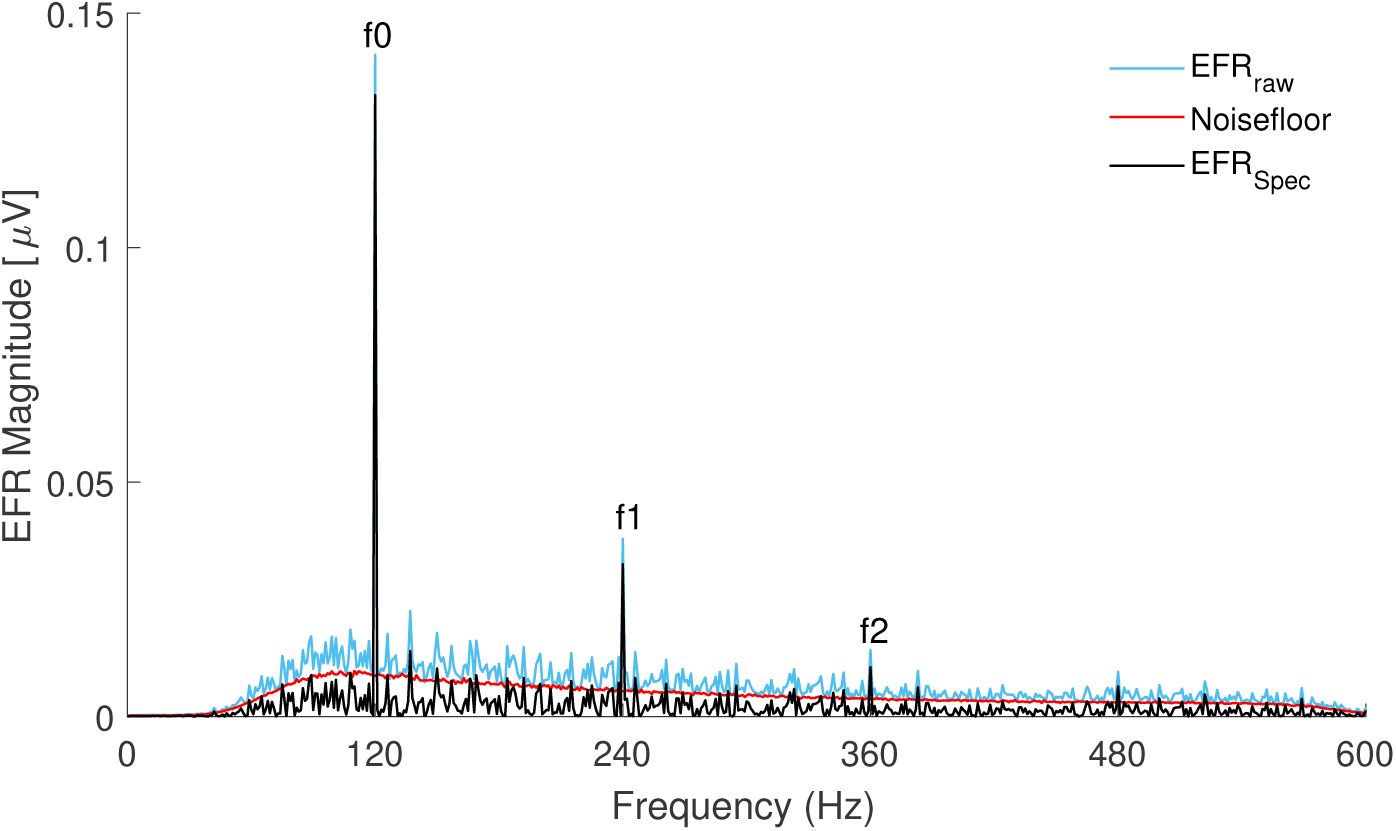
Magnitude spectrum of the EFR_raw_(f) (in blue), Noisefloor(f) (in red) and EFR_Spec_(f) (in black) calculated for subject No. 8 from the first experiment. EFR spectra were evoked by the stimulus with the broadest bandwidth, i.e. [0.25-22] kHz. Peaks at the stimulus modulation frequency, and two harmonics (i.e. f_0_ = 120Hz, f_1_ = 240Hz and f_2_ = 360Hz) are clearly visible above the noise-floor.

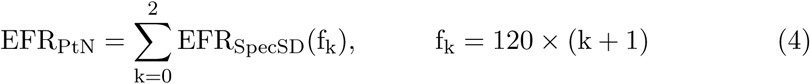

To construct DBEFRs, the calculated EFR_PtN_ for each narrower-band condition was subtracted from the following wider-band condition using: 

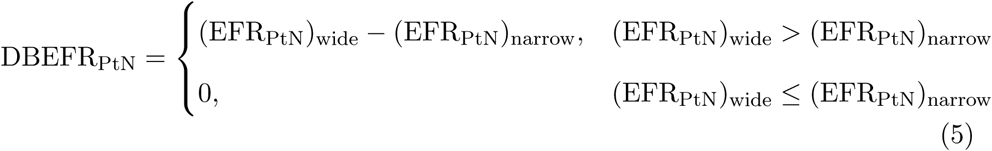

Derived frequency bands from EFRs to the first experimental stimuli are shown schematically in Fig. 2b.

## 4. Questionnaire analysis

The completed questionnaires from the participants in the first experiment were used to estimate the individual life-time noise exposure dose. To this end, the collected individual data related to the frequency and duration of experienced noise exposure were converted to a number of sessions per year multiplied by the duration and the personal estimated noise loudness scores, i.e. a number between 1 and 5. We followed the procedures as described in Degeest et al. (2014). The scores were separately calculated for questionnaire categories: (i) playing musical instrument in a band, (ii) attending festivals, concerts and discotheques and (iii) using noisy tools. Outcomes were normalized across NH and NHSR groups participants by the highest reported dose, i.e. 30600, 18480 and 26000 hours in each category, respectively.

## 5. Model Simulations

A biophysical model of the human auditory periphery (Verhulst et al., 2018a), schematically shown in Fig. 4, was adopted to simulate the experimental conditions and to investigate the effect of different aspects of sensorineural hearing deficits on the EFR_PtN_ and DBEFR_PtN_ magnitudes. The original implementation of the model is described in Verhulst et al. (2018a) and can be downloaded from *“https://github.com/HearingTechnology/Verhulstetal2018Model“*. The parameters which determine the weights between the population AN, cochlear nucleus (CN) and inferior colliculus (IC) responses were adjusted along with the AN innervation patterns across CF for the purpose of this study.

**Figure 4.**
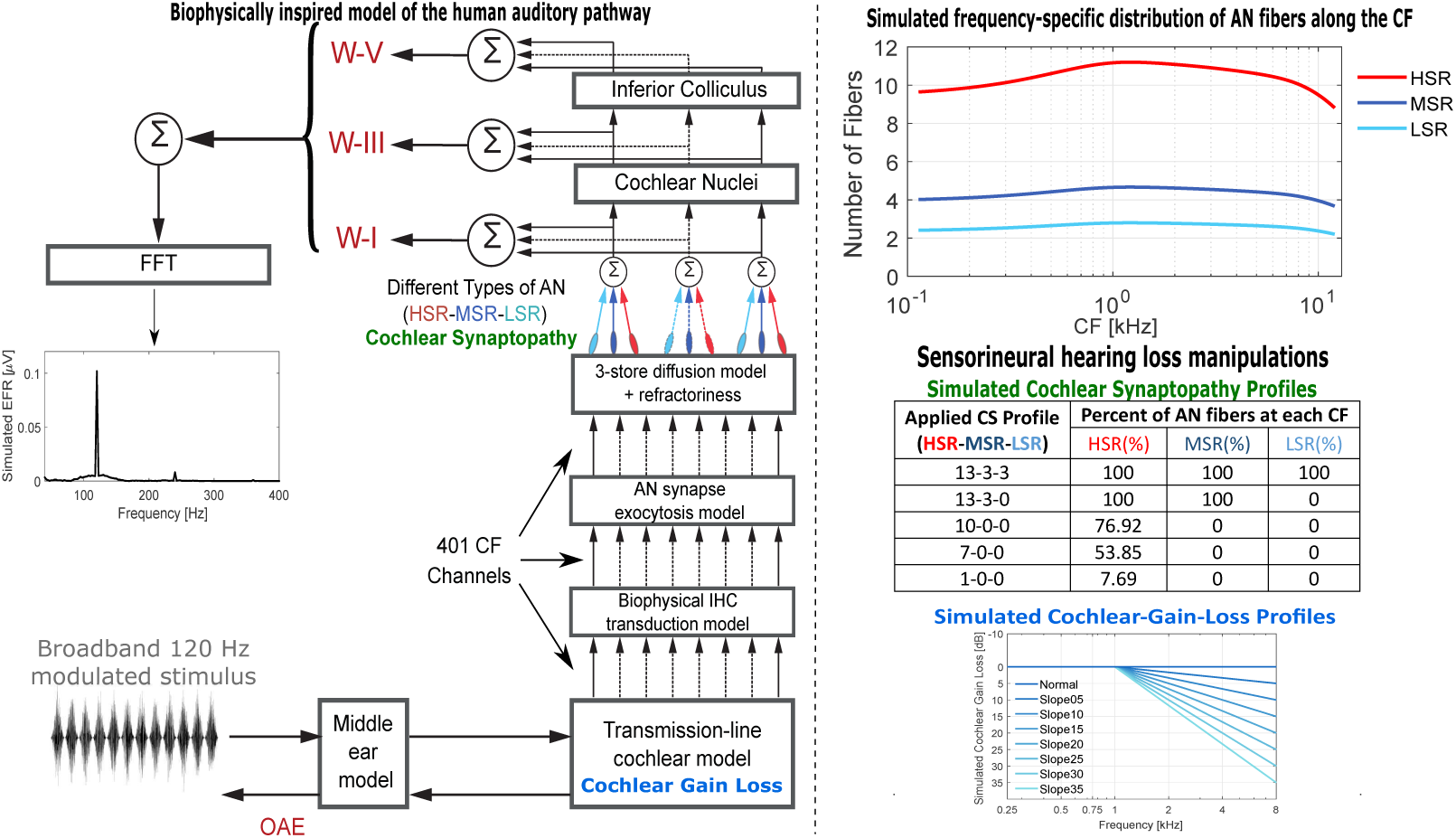
Modeling approach. The block-diagram shows different levels of the auditory pathway modelled in the employed biophysical model of the hearing periphery (Verhulst et al., 2018a). The top-right graph indicates the simulated distribution of different types of AN fibers across CF. The table shows simulated CS profiles and the graph on the bottom right depicts simulated different degrees of cochlear gain loss. The corresponding simulated thresholds at 8 kHz are indicated by the legend.

### 5.1. Auditory nerve-fiber distribution

The original model implementation introduced the same number of synapses between inner-hair-cells (IHCs) and AN fibers for all simulated characteristic frequencies (CF), whereas human and rhesus monkey innervation patterns show a bell-shaped pattern across CF. To make the model more realistic, the averaged synaptic counts of four control rhesus monkeys (seven ears) and nine frequencies (Valero et al., 2017) were mapped to corresponding fractional distances of the human cochlea using the monkey place-frequency map (Greenwood, 1990). Fractional distances from the base of cochlea, d_i_, were calculated according to the measured frequency points 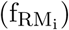: 

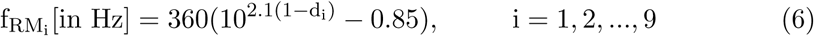

The obtained d_i_s were substituted into the analogous Greenwood map equation for humans, yielding the corresponding frequency points 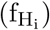: 

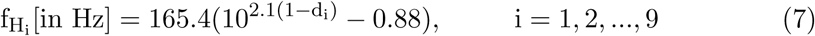

To calibrate the model with the applied AN pattern, a 70 dB-nHL click-train containing both stimulus polarities was presented at a rate of 11 Hz. To perform this calibration, simulated ABR wave amplitudes were matched to the experimental data on the basis of 55 averages. Specifically, the M_1_ = 4.6729 *×* 10^−14^, M_3_ = 5.6885 *×* 10^−14^ and M_5_ = 14.641 *×* 10^−14^ parameters were adjusted on the basis of average NH ABR wave-I, III and V reference data from Picton (2010), i.e. w_I_ = 0.15*µ*V_p_, w_III_ = 0.17*µ*V_p_ and w_V_ = 0.61*µ*V_pp_.

Using the synapse counts from rhesus monkey and the mapped frequency points for the human cochlea 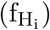, a “smoothing spline” curve was fit to estimate the number of synapses across all frequency channels in the model. Finally, to simulate different AN fiber types, i.e. high spontaneous-rate (HSR), medium spontaneous-rate (MSR) and LSR fibers, and their properties, the obtained population distribution was multiplied by the corresponding AN type proportion factor C, i.e. C_HSR_ = 0.60, C_MSR_ = 0.25 and C_LSR_ = 0.15 (Liberman, 1978, cat data), before responses were summed at each simulated CF and fed to the CN model. The simulated frequency-specific AN fibers distribution is shown on the top-right column of Fig. 4.

### 5.2. Stimuli

The model stimuli were matched to the experimental conditions and had a duration of 600 and 400 ms for the first and second experiment, respectively. Twenty stimulus repetitions with different white noise iterations were applied to the model and simulations were averaged before the EFR_PtN_ was calculated using the same procedure as in Eq. 4. The amplitudes of the model stimuli were set based on the broadest condition, i.e. 0.25 to 22 kHz for the first experiment and 0.3 to 16 kHz for the second experiment to yield an input of 70 dB SPL. The narrower band stimuli were calibrated relative to the broadest condition, such that they had the same spectral level as the broadband condition but with a different SPL.

### 5.3. Simulating sensorineural hearing loss

The simulated CS profiles and their corresponding AN fiber types are shown in Fig. 4. Different degrees of CS were modelled by manipulating the number and types of AN fibers. The table in Fig. 4 shows the simulated synaptopathy profiles. OHC damage was simulated by changing the CF-dependent mechanical gain of the cochlea by moving poles of the BM admittance function to yield a filter gain reduction corresponding to a desired dB-HL-loss, which also yielded wider cochlear filters. The inset in Fig. 4 shows the simulated cochlear gain loss profiles. Procedures are further detailed in Verhulst et al. (2016, 2018a).

## 6. Results

### 6.1. EFR and dependence on stimulus frequency

Figure 5 shows individual and group-mean EFR_PtN_ magnitudes to different frequency bandwidths in the first (panel a) and second (panel b) experiments. Despite within-group individual variability, experimental group-means revealed approximately constant EFR_PtN_ magnitudes to stimuli with frequencies below 2 kHz and reduced magnitudes to frequencies above 2 kHz and 2.8 kHz in the first and second experiment, respectively. A paired-sample t-test with Bonferroni correction was applied to compare EFR_PtN_ magnitudes to stimuli with different frequency bandwidths in each group. In the first experiment, a single significant difference was observed between the EFR_[2−22]_ and EFR_[4−22]_ conditions in NH group (t(11)=7.02, p*<*0.0000; specificed by # in Fig. 5a), which disappeared for the NHSR group (t(8)=3.13, p=0.014). In the second experiment, a paired-sample t-test with Bonferroni correction gave a significant difference between EFR_[2.8−16]_ and EFR_[5.6−16]_ in yNH (t(12)=7.86, p*<*0.0000; specificed by + in Fig. 5b) and oNH groups (t(12)=6.21, p*<*0.0000; specificed by ++ in Fig. 5b), but not in the oHI group (t(9)=2.03, p=0.072). Simulated NH-EFRs are shown in hexagons in Fig. 5 and corroborate experimental findings by showing a minor contribution of stimulus frequencies below 2 kHz on the EFR generation.

**Figure 5.**
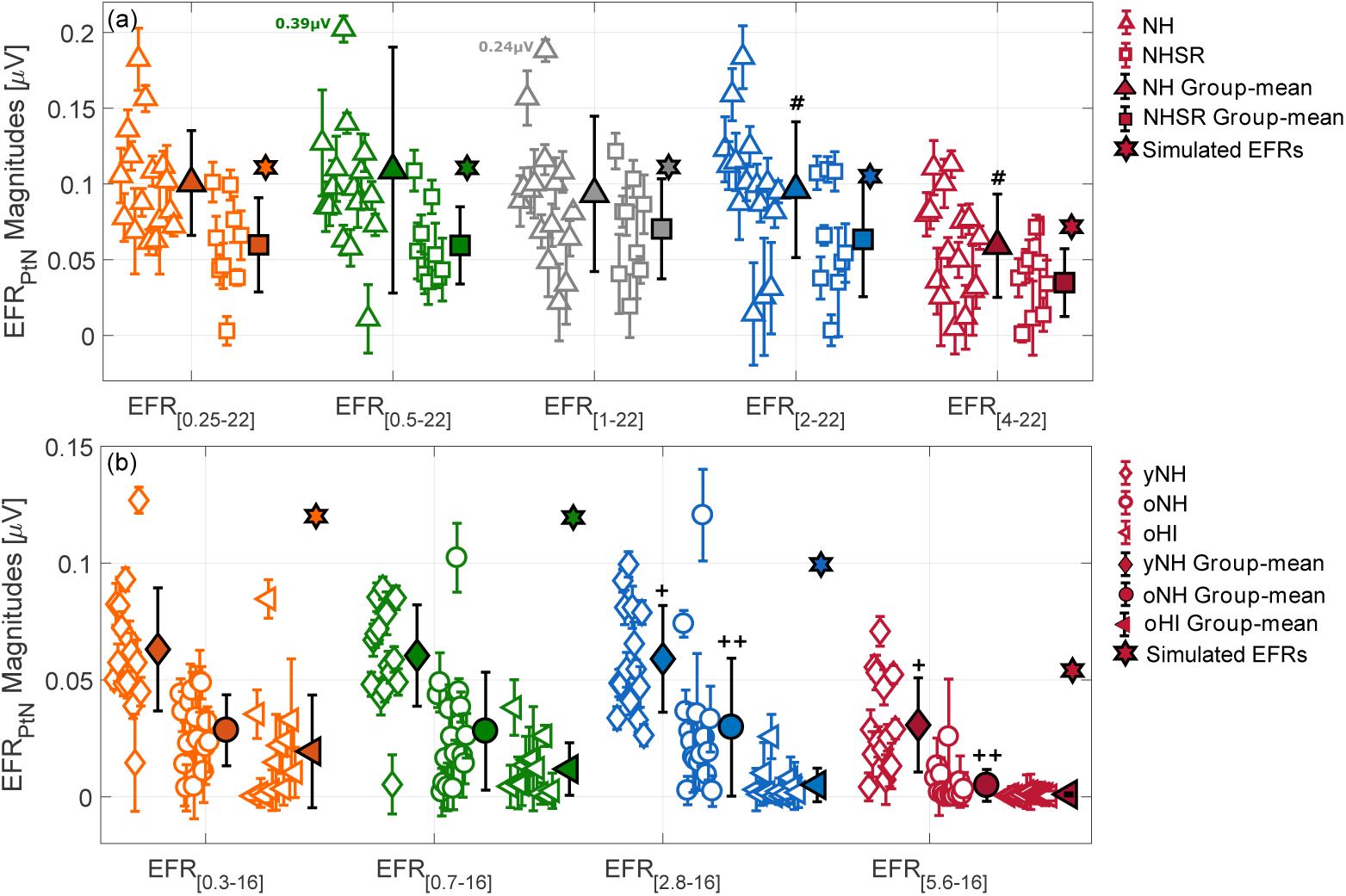
EFR_PtN_ magnitudes to 120-Hz modulated stimuli with different white noise carrier bandwidths in the (a) first and (b) second experiment. Individual data-points are depicted with open symbols and standard deviations were obtained using a bootstrapping procedure (Zhu et al., 2013). Filled symbols reflect the group-means and their corresponding standard deviations. Simulated EFRs from a NH model were added in filled hexagons. Significant effects of considered frequency-band on EFR_PtN_ magnitudes are specified by: (#) in the NH-group (first experiment), (+) in the yNH-group and (++) in the oNH-group (second experiment). To enhance the visualization of differences, panel (a) was plotted on narrower y-axis range, therefore the real values of lowered EFR_PtN_ magnitudes were specified next to the corresponding data-points.

### 6.2. Derived-Band Envelope Following Responses (DBEFRs)

DBEFR_PtN_ magnitudes calculated using Eq. 5 are shown in Fig. 6 for the first (panel a) and second (panel b) experiment. A paired-sample t-test with Bonferroni correction comparing the DBEFR_PtN_ magnitudes in each group revealed only a significant difference between the [1-2] and [2-4] kHz condition in the NH group (t(11)=-3.99, p=0.002; specificed by # in Fig. 6a). In the second experiment, paired-sample t-test showed significant difference between [0.3-0.7] and [2.8-5.6]-kHz conditions only in yNH group (t(12)=-7.00, p*<*0.000; specificed by + in Fig. 6b). In support of our experimental findings, simulated NH-DBEFR magnitudes in both experiments (shown by hexagons in Fig. 6a and b) were equal for derived-bands below 2-kHz and increased for DBEFR_[2−4]_ (in the first experiment) and DBEFR_[2.8−5.6]_ (in the second experiment). In line with EFR_PtN_ findings in Section 6.1, experimental and simulated DBEFR_PtN_ magnitudes in both experiments showed an increased contribution of the [2-6] kHz derived frequency band to the EFR generation.

**Figure 6.**
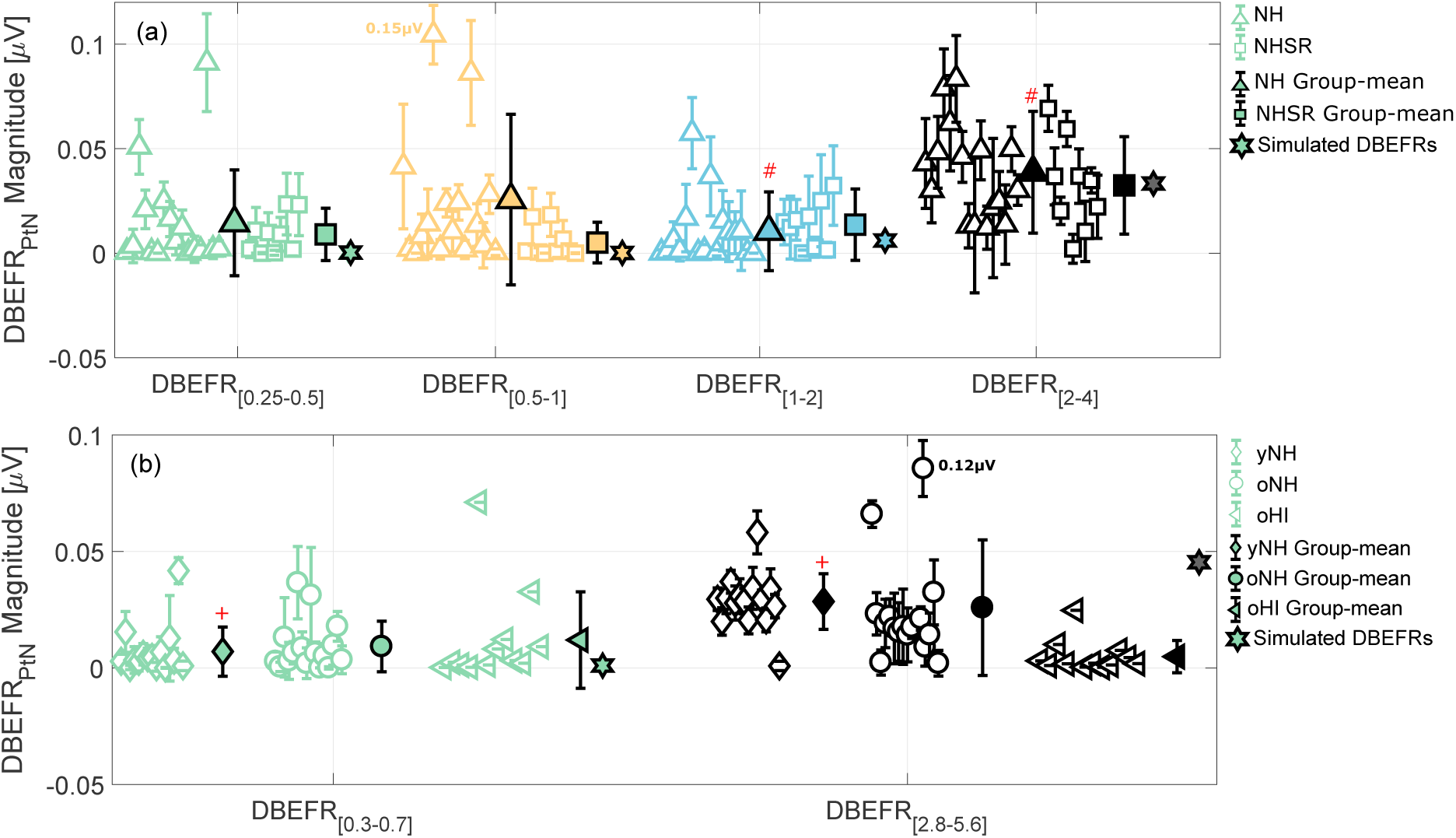
DBEFR_PtN_ magnitudes derived using Eq. 5 for 120 Hz modulated stimuli with different white-noise-carrier bandwidths in the (a) first and (b) second experiment. DBEFR_PtN_ for each frequency band was obtained from a wider and narrower width stimulus. Standard deviations were calculated using a bootstrapping procedure and stemmed from averaged responses from 20 stimulus iterations in the model simulations. Group means and standard deviations are depicted using filled symbols. Significant effects of considered frequency-band on NH-group in the first experiment and yNH-group in the second experiment are specified by (#) and (+), respectively. To enhance the visualization of differences, figures were plotted on narrower y-axis range, therefore the real values of lowered DBEFR_PtN_ magnitudes were specified next to the corresponding data-points.

### 6.3. Possible origins of individual EFR differences

Previous studies have shown a dependency of the scalp-recorded AEP magnitude to head size, sex and age (Trune et al., 1988; Mitchell et al., 1989; Vasilkov and Verhulst, 2019, preprint). Hence, the spread of data-points within different recorded test-groups and spectral bandwidths could be explained by subject-specific factors unrelated to hearing or hearing-related factors associated with the main factors for grouping: (i) self-reported hearing difficulties in noisy environments in the first experiment, (ii) age and (iii) elevated hearing thresholds in the second experiment.

Pooling together the NH and NHSR EFR_PtN_ magnitudes, a regression analysis was conducted to investigate the effect of age, 4 kHz threshold, head size and DPTH_3000_ on the EFR_[2−22]_ (Fig. 7, left column) and DBEFR_[2−4]_ magnitude (Fig. 8, left column). None of the regressions showed a relation between tested variables, suggesting that other factors than those reported were responsible for the individual variability among listeners. The regression analysis on EFR_PtN_ and DBEFR_PtN_ magnitudes combined from all experimental groups in the second experiment (Fig. 7 and 8, right column) showed a meaningful correlation of age, threshold, head size and DPTH_4000_ with the EFR_[2.8−16]_ magnitude. However, extracting the DBEFR_[2.8−5.6]_, reduced the correlation with age and 4-kHz threshold and suppressed any meaningful correlation with head-size and DPTH_4000_. More-over, excluding the oHI group from the correlation analysis, led to a reduced and insignificant correlation coefficient (R=-0.382, p=0.083) between 4-kHz threshold and DBEFR_[2.8−5.6]_. These results suggest that the proposed DBEFR metric is not affected by head size. Moreover, individual variabilities between the yNH and oNH groups in the second experiment might be related to degraded temporal envelope coding as a consequence of CS (Bharadwaj et al., 2015), given the insignificant correlations of DBEFRs with the 4-kHz threshold, DPTH_4000_ and head size.

**Figure 7.**
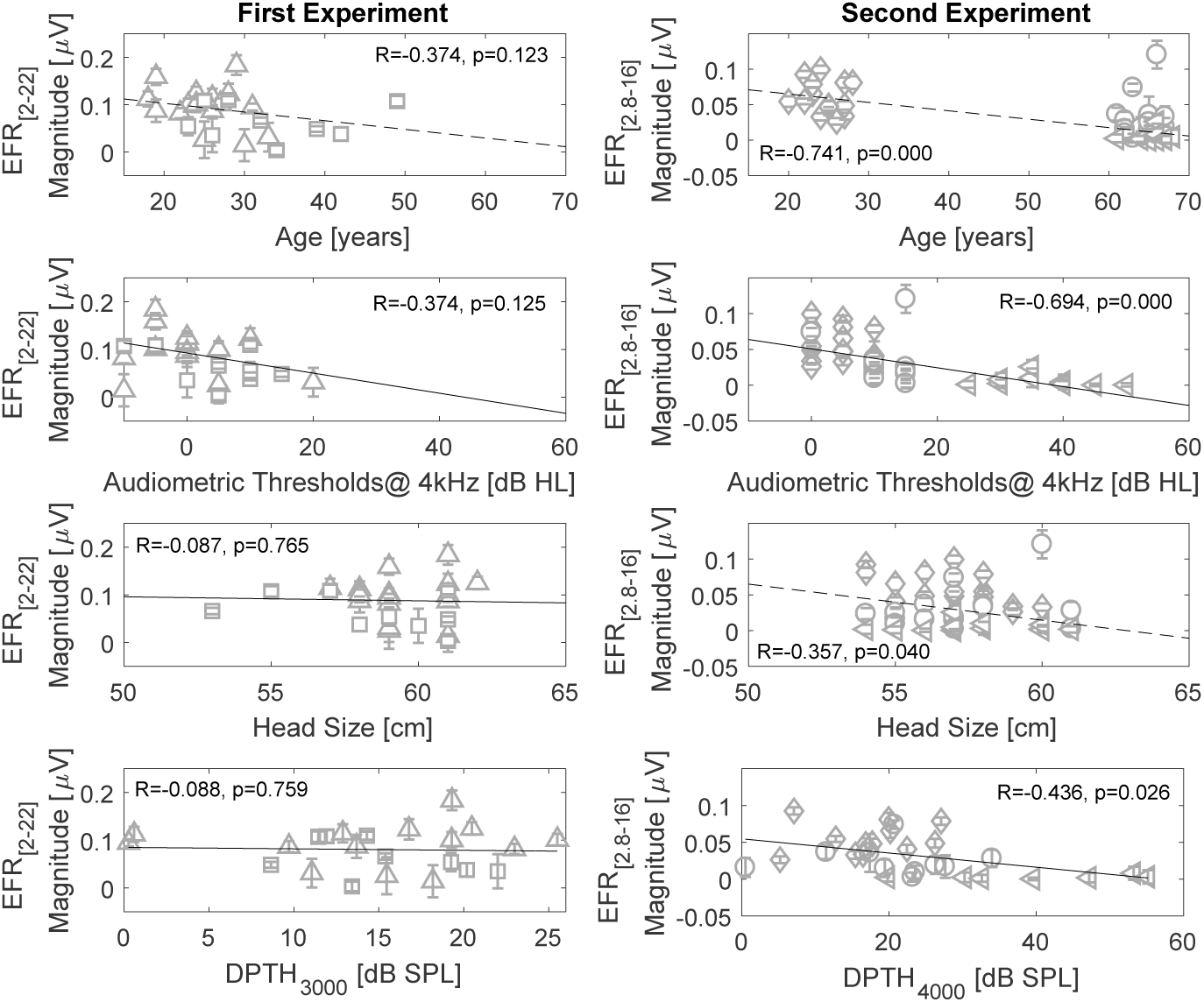
Correlation analysis of EFR_[2−22]_ (EFR_[2.8−16]_) with age, audiometric threshold at 4 kHz, head-size and DPTH_3000_ (DPTH_4000_) in the first (left) and second (right) experiments. Correlation between EFR magnitudes and all factors but age were reported using the Pearson’s correlation coefficient. The Spearman’s correlation coefficient was calculated to study the effect of age in the second experiment.

**Figure 8.**
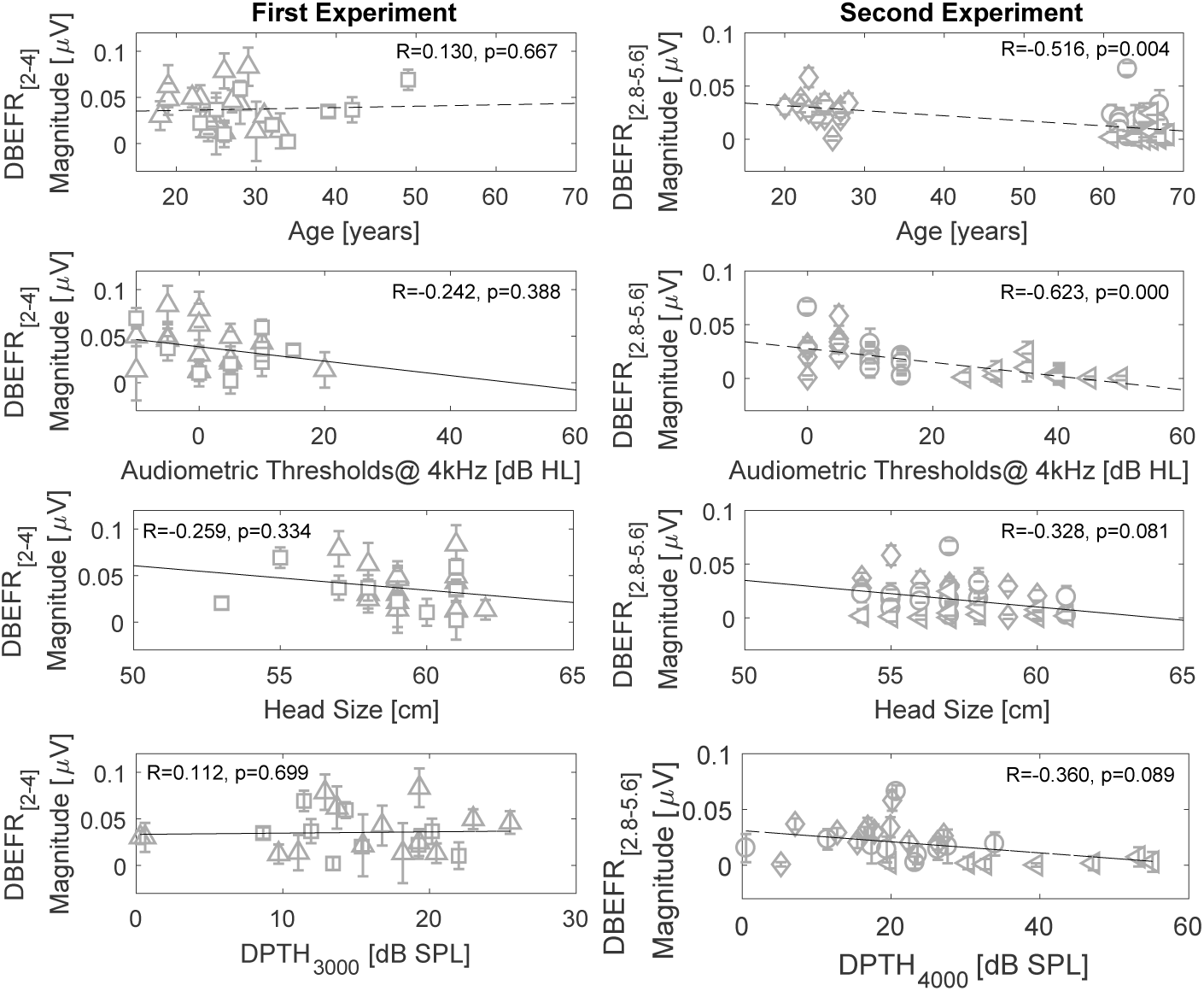
Correlation analysis of DBEFR_[2−4]_ (DBEFR_[2.8−5.6]_) with age, audiometric threshold at 4 kHz, head-size and DPTH_3000_ (DPTH_4000_) in the first (left) and second (right) experiments. Correlation between DBEFR magnitudes and all factors but age were reported using the Pearson’s correlation coefficient. The Spearman’s correlation coefficient was calculated to study the effect of age in the second experiment.

### 6.4. EFR_PtN_ and DBEFR_PtN_ magnitude variability across tested groups

To investigate the separability of the recruited groups by means of their DBEFR magnitudes, we analysed the group-mean differences in each experiment. In the first experiment, an independent two-sample t-test comparison between the means of stimulated frequency bandwidths in the NH and NHSR group (Fig. 5a), showed a significant difference only between the [2-22] and [4-22]-kHz conditions (EFR_[2−22]_: t(19)=3.36, p=0.003 and EFR_[4−22]_: t(19)=2.76, p=0.012). However, significant mean-differences disappeared between similar conditions in the NH and NHSR groups after extracting DBEFR magnitudes in Fig. 6a (DBEFR_[2−4]_: t(19)=0.90, p=0.338). The insignificant difference across groups and insignificant correlation coefficients of DBEFR_[2−4]_ with subject-specific factors observed in Fig. 8, might partly be explained by the different amounts of experienced lifetime noise exposure reported in the questionnaires and might point to various degrees of noise-induced CS. Calculated noise scores in Fig. 9 revealed an insignificant correlation with DBEFR_[2−4]_ magnitudes (R=0.13, p=0.089). However, certain cases appeared to be inconsistent with our noise-induced synaptopathy hypothesis, i.e., (i) high noise scores in the NH group, e.g. subject No. 12 and (ii) low noise scores in the NHSR group, e.g. subject No. 1. We suggest that the insignificant group-mean differences can be explained by (i) subject-dependent unreliable discriminating factor between NH and NHSR group (Coughlin, 1990), (ii) variability in answering lifetime noise-exposure dose in questionnares (Prendergast et al., 2017; Bramhall et al., 2017), (iii) an insufficient number of samples and (iv) a limited sensitivity of the DBEFR_PtN_ metric to noise-induced CS.

**Figure 9.**
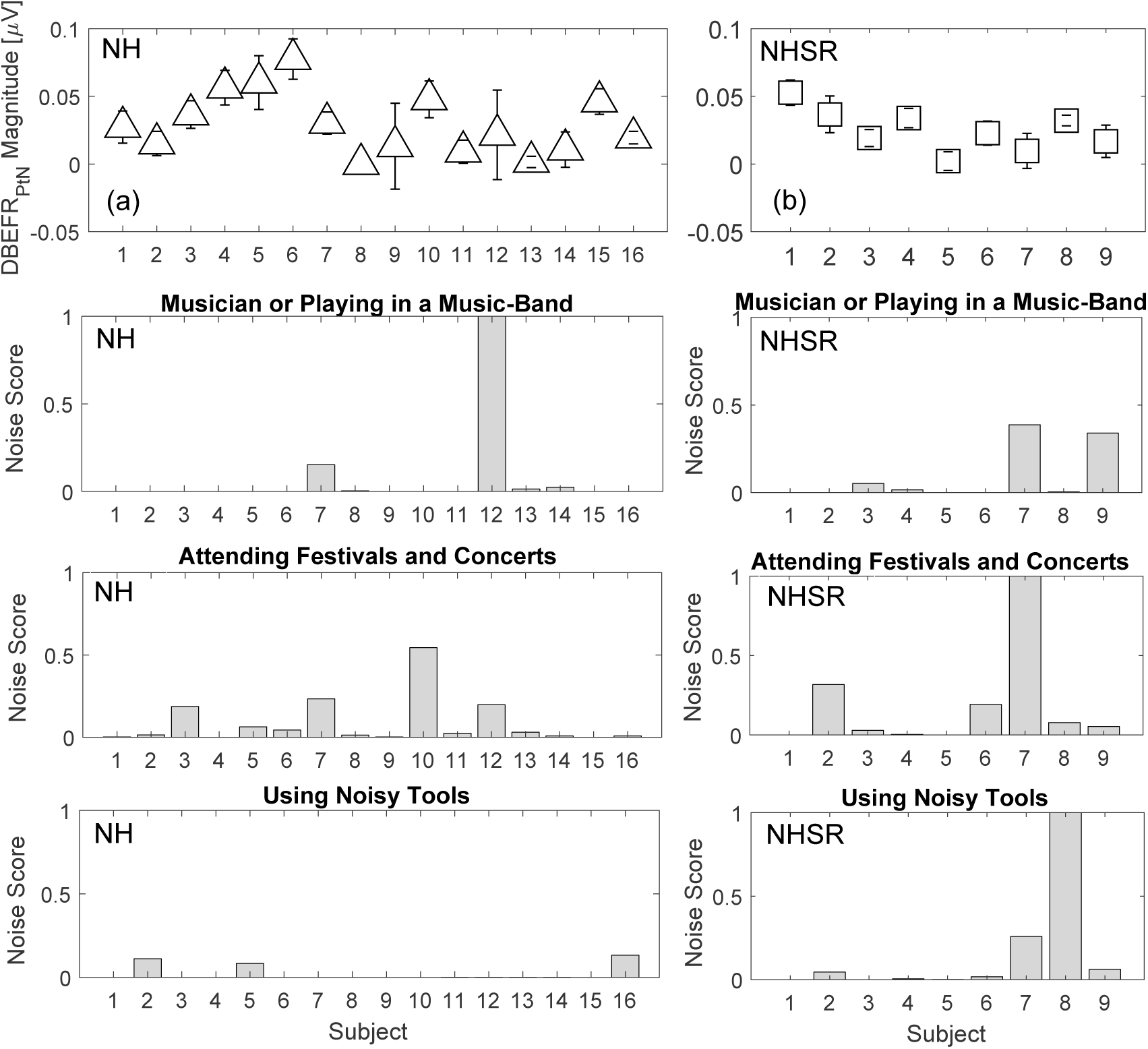
Bar-plots of noise scores acquired from questionnaires of NH and NHSR groups, classified in three categories, i.e. experience noise as a consequence of (i) playing a musical instrument in a band, (ii) attending festivals or concerts and (iii) using noisy tools. Results are shown normalised, where the score of 1 corresponds to 30600, 18480 and 26000 hours of accumulated noise dose on the considered categories, respectively.

In the second experiment, an independent two-sample t-test was applied to investigate the effect of age between the yNH and oNH groups, and elevated high-frequency thresholds between the oNH and oHI groups. This comparison showed a significant effect of age on all frequency bandwidths and a significant effect of hearing threshold on all frequency bands except for the [5.6-16] kHz band (t(21) = −1.81, p = 0.084). The same comparison for the DBEFR magnitudes revealed a significant effect of age and hearing threshold only in the [2.8-5.6]-kHz derived band condition (t(24) = 3.13, p=0.004 and t(21) = −4.60, p = 0.002, respectively), consistent with the correlation presented in Fig. 8. Detailed *t* and *p* values of independent two-sample t-tests, evaluating the effect of age and hearing thresholds on EFR and DBEFR magnitudes, are listed in Table. 1.

**Table 1:**
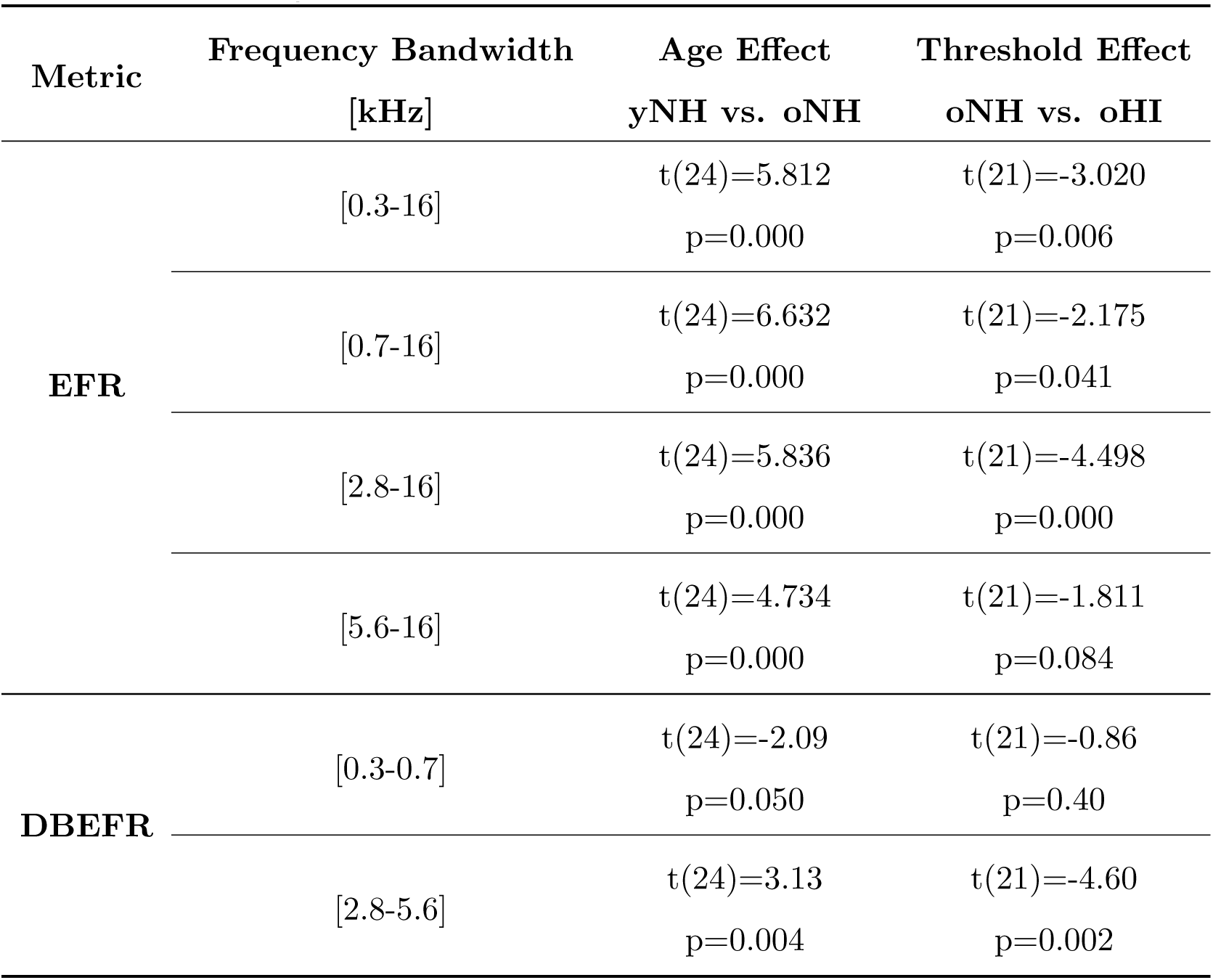
The results of a two-tailed *t* test show the effect of age and hearing threshold on EFR and DBEFR magnitudes in the second experiment.

Our group-mean results combined with the correlation analysis in Section 6.3 suggests that the DBEFR metric removes inter-subject variability unrelated to hearing between yNH and oNH groups, but leaves individual magnitude differences within a group meaningful, given the often non-overlapping standard deviations. Consequently, the significant group-mean difference between yNH and oNH might reflect individual degrees of sensorineural hearing loss. To investigate the diagnostic sensitivity, it is of course necessary to understand the respective role of OHC deficits and CS on DBEFR magnitudes. Given that oHI listeners may suffer from both OHC deficits and CS, it is important to study the impact of OHC-damage and CS, both independently and concomitantly.

### 6.5. The EFR relationship to different aspects of sensory hearing-loss

Since OHC-damage and CS might both affect the EFR magnitude (Garrett and Verhulst, 2019; Vasilkov and Verhulst, 2019, preprint), we employed a computational model of the auditory periphery to simulate how different degrees of CS affected the EFR_PtN_ magnitude, both in presence and absence of high-frequency sloping OHC-loss above 1 kHz (simulated high-frequency sloping audiograms in Fig. 4). The most sensitive regions of the cochlea responding to a 120-Hz modulated broadband noise were identified to lie between the CFs of 2 and 6 kHz (Keshishzadeh et al., 2019). As a result, we only considered two EFR conditions of each experiment, namely EFR_[2−22]_ and EFR_[4−22]_ in the first experiment (Fig. 10a) and EFR_[2.8−16]_ and EFR_[5.6−16]_ in the second experiment (Fig. 10b). Model simulations showed that CS, when no other hearing deficits co-occur, reduces the EFR and DBEFR magnitudes. Applying sloping high-frequency OHC-damage increased the DBEFR magnitudes in both experiments (Fig. 10c and d). According to the simulations, the NH DBEFR magnitude reduced by 46% as a consequence of removing 47% of the AN fibers (i.e., the 10-0-0 CS profile defined in Fig. 4), while the Slope20 OHC-damage (defined in Fig. 4) increased the NH DBEFR magnitude by 27%. Hence, the effect of OHC-damage on the DBEFR magnitude is smaller than that of CS alone, however it is not negligible. Therefore, the experimental range of individual EFR and DBEFR magnitudes can be explained by different degrees of variation simulated by CS and OHC-damage.

**Figure 10.**
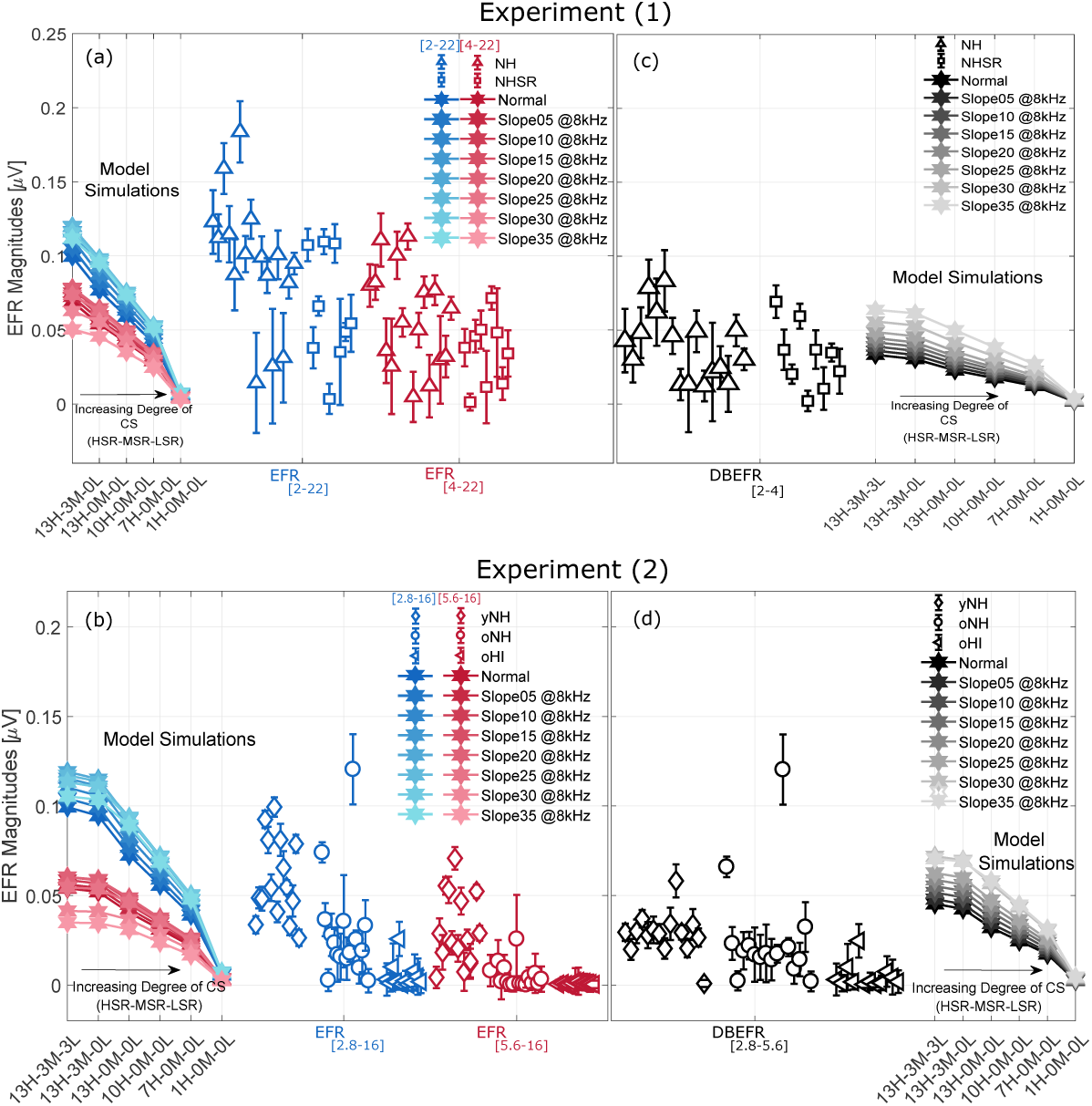
Experimental EFR_PtN_ and DBEFR_PtN_ magnitudes (colored open symbols): (a) EFR_PtN_ to [2-22] and [4-22] kHz, (b) EFR_PtN_ to [2.8-16] and [5.6-16] kHz and (c) DBEFR_PtN_ at [2-4] kHz and (d) DBEFR_PtN_ at [2.8-5.6] kHz. Simulated EFR_PtN_ (a,b) and DBEFR_PtN_ (c,d) magnitudes are shown in each panel using filled hexagons and degrees of CS as indicated on the X axis and CF-dependent patterns of OHC damage as given by the legend.

Our simulations predicted the experimental observed absolute range of DBEFR magnitudes and explained the experimental differences between yNH and oNH groups on the basis of age-induced CS, not OHC-damage induced differences. Furthermore, the simulations suggest that oNH and oHI listeners might both suffer from CS. Results are less clear for the NHSR group where there is a strong overlap with the NH group. However, the noise scores from the questionnaires in Fig. 9, could ascribe some of the spread in DBEFR magnitudes within the NH and NHSR groups to noise-induced CS, and to a lesser degree to OHC-damage given all had normal hearing thresholds.

It is worthwhile to note that EFR magnitudes in both experiments (Fig. 10a and b), decreased as a result of CS alone and increased by applying high-frequency OHC-damage with a severity of less than 20 dB-HL at 8 kHz. However, higher degrees of OHC-damage reduced the EFR magnitudes. We explain this non-monotonic behaviour on the basis of the AN fiber discharge rate-level curve, where increased simulated EFR_PtN_ magnitudes (Fig. 10 c and d) and amplitude-modulated (AM) responses (Fig. 11b) to supra-threshold stimuli (70 dB-SPL) caused by OHC-damage, might stem from the extended dynamic range of the AN fibers for less effective AN-driving levels (Bharadwaj et al., 2014, their Fig. 3c). Given that experimental and simulated stimuli were calibrated to have equal spectral magnitudes for all stimulus bandwidths, the narrowest bandwidth stimulus was presented at a lower overall sound level than the 70 dB-SPL broadband stimulus. Thus, applying more severe OHC-loss, lowered the AN discharge rate and envelope synchrony strength (Verhulst et al., 2018a, Fig. 5) and decreased the EFR magnitudes (Verhulst et al., 2018a, their Fig. 7). However, DBEFR magnitudes increased monotonically for all simulated degrees of OHC damage (Fig. 10c and d).

**Figure 11.**
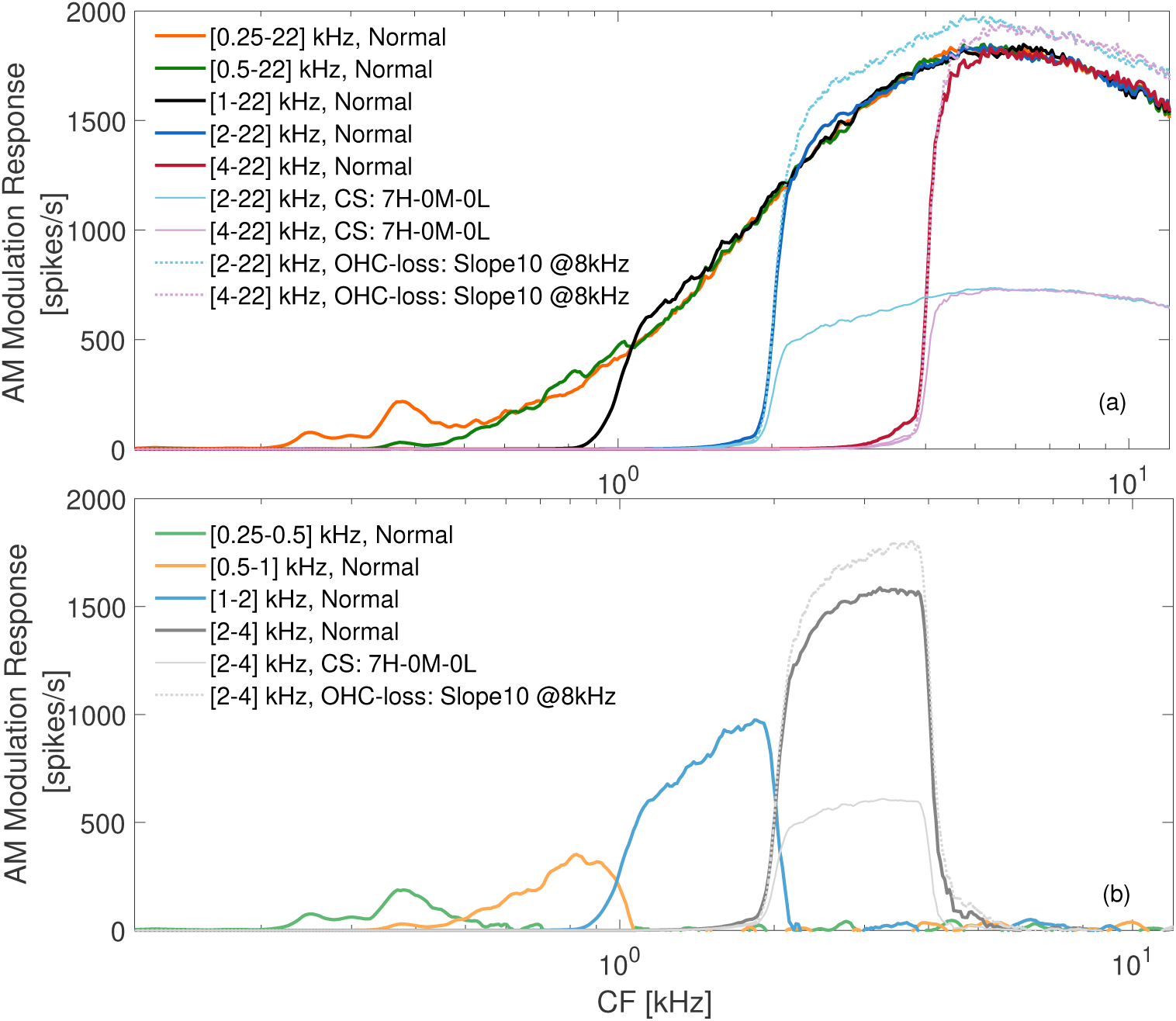
Modulated responses calculated at each CF using Eq. 8 and 9 to different experimental conditions for normal listeners and different sensorineural hearing losses at the AN processing level of the model, (a) broad-band and (b) derived-band. In both panels, dotted lines show AM-responses to sloping 10 dB-HL OHC-loss at 8 kHz and lighter colors indicate AM responses to certain degree of CS.

## 7. Discussion

### 7.1. Tonotopic sensitivity of the EFR generators

Despite the individual variability within groups, experimental group-mean EFR_PtN_ magnitudes to broadband stimuli with different bandwidths (Fig. 5a), were equal at frequencies below 4 kHz and reduced in response to [4-22] kHz condition. In the second experiment (Fig. 5b), the EFRs remained equal at frequencies below 5.6 kHz and degraded when the [5.6-16] kHz band was added. Consequently, equal DBEFR_PtN_ magnitudes were obtained for frequencies below 2 kHz. Individual variability was best observed for the DBEFR_PtN_ extracted from the [2-4] kHz (first experiment, Fig. 6a) and [2.8-5.6] kHz (second experiment, Fig. 6b) frequency bands. Simulated EFRs to the experimental stimuli shown with hexagons in Fig. 5 and 6, confirmed observed experimental EFR_PtN_ and DBEFR_PtN_ frequency-dependent behaviour. In addition, the model can be used to study which CF regions along the cochlea contributed strongly to the population EFR response. To this end, we calculated the AM (Fig. 11a) and derived-band AM (DBAM) responses at each CF (Fig. 11b) as follows: 

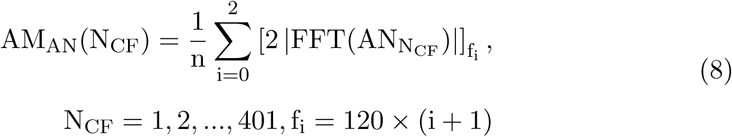

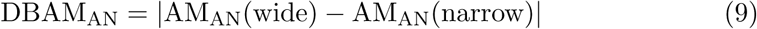

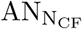 is the AN-response at N_CF_ channel and n = n_1_ as was defined in Eq. 1. These simulations corroborate the experimentally-observed minor contribution of low-frequency CF channels to the EFR generation.

In a previous modelling study (Keshishzadeh et al., 2019), we investigated the tonotopic sensitivity of EFR_PtN_ to broadband stimuli and ascribed the poor low-frequency AM coding to a combination of the chosen modulation frequency (120 Hz) and the narrower bandwidth of apical cochlear filters compared to the higher CF filters (Moore and Glasberg, 1983). Model simulations in response to the spectrally broadest condition, i.e. [0.25-22] kHz, modulated with a range of lower modulation frequencies than 120 Hz, showed that the saturation properties of AN fibers limited the modulation response at all modulation frequencies at higher CFs despite an enhanced modulated response at the BM. This resulted in a degraded response at CFs above 4 kHz and shifted the frequency sensitivity of AM coding to the lower CFs at low modulation frequencies. Since the brain response to modulation frequencies below 70 Hz may contain cortical as well as brainstem contribution (Purcell et al., 2004; Picton, 2010, Chapter 10), employing low modulation frequencies might render EFR-based CS diagnosis insensitive, even though an improved frequency-sensitivity can be obtained from the apical regions using these lower modulation frequencies. Therefore, the employed experimental modulation frequency, i.e. 120-Hz in combination with a broadband carrier, might be able to establish a frequency-specific CS diagnosis at frequencies above 2 kHz. In this context, the proposed DBEFR method showed a notable contribution of the [2-4] kHz CF region to the EFR generation by showing a significantly stronger DBEFR_PtN_ magnitude compared to lower derived-band conditions in the NH group.

### 7.2. Diagnostic Applications

The measured DBEFR magnitudes are individually separable and above the noise-floor even for HI listeners, whose group-mean was significantly above the noise-floor. In addition, the DBEFR offers a frequency-specific metric to assess supra-threshold temporal coding of the population of AN fibers and brainstem neurons in the [2-6] kHz region. Despite these promising results, the diagnostic sensitivity of DBEFRs also has limitations. The proposed DBEFR magnitude is sensitive to CS alone, when no other coexisting hearing deficits occur and is hence applicable for use in ageing listeners with normal audiograms and those with self-reported hearing difficulties or prone to noise exposure. However, DBEFRs are also affected by OHC damage (Fig. 10). The metric hence needs to be complemented with another supra-threshold metric sensitive to OHC damage within the tonotopic range of interest to allow a separation of the CS and OHC aspect of sensorineural hearing damage from the recorded DBEFRs from listeners with impaired audiograms.

Lastly, the employed high modulation frequency, i.e. 120 Hz, suppresses cortical contributions to the EFR_PtN_ magnitudes, but also degrades AM-coding from lower CFs and thereby limits the tonotopic sensitivity of the EFR_PtN_ to frequencies above 2 kHz. Consequently, apical-end supra-threshold hearing deficits would not be reflected in the proposed DBEFR_PtN_ metric even for stimuli which contain frequencies below 2 kHz. These results are consistent with the source generators of derived-band ABRs (DBABR), which reduce in amplitude for bands below 2 kHz (Don and Eggermont, 1978). This predominant basal origin of the ABR also confines the potential of ABR/DBABR-based CS diagnosis to basal cochlear regions (e.g. wave-I amplitude).

## 8. Conclusion

We proposed the use of a relative DBEFR_PtN_ metric to render the EFR_PtN_ frequency-specific and rule out subject-specific factors unrelated to hearing to apply it in the study of identifying the origins of sensorineural hearing deficits and clarifying their role in supra-threshold temporal envelope encoding. DBEFR_PtN_ magnitudes from two experiments were analysed and compared to model simulations to conclude that the frequency-sensitivity of DBEFR_PtN_ magnitudes to broadband stimuli is limited to the [2-6] kHz bandwidth. Secondly, we showed that the DBEFR metric eliminates inter-subject variability caused by hearing-unrelated sources. Model simulations (Fig. 10) explained the significant difference between yNH and oNH listeners on the basis of CS, which could result from age-induced CS as identified from human post-mortem studies (Makary et al., 2011; Viana et al., 2015; Wu et al., 2019). Supported by model predictions (Fig. 10d), the significant difference between age-matched oNH and oHI groups was explained by OHC-damage and coexisting CS as a consequence of ageing. Accordingly, profound OHC damage may confound DBEFR-based clinical applications of CS diagnosis. Despite this limitatiion in the differential diagnosis of CS and OHC deficits on the basis of the DBEFR magnitude, the proposed metric can be used to diagnose CS in a frequency-specific manner in listeners with thresholds below 20 dB-HL. Moreover, it provides an objective marker of supra-threshold temporal envelope coding, which can be used to study its role in sound perception studies. Lastly, our results clearly demonstrate that older listeners with or without impaired audiograms suffer from degraded temporal envelope coding at frequencies above 2 kHz.

## Acknowledgement

This work was supported by European Research Counsil (ERC) under the Horizon 2020 Research and Innovation Programme (grant agreement No. 678120 RobSpear).

